# IL27 exerts a powerful effect on systolic overload-induced cardiac inflammation, fibrosis, and heart failure development

**DOI:** 10.64898/2026.08.05.743140

**Authors:** Ziru Niu, Umesh Bhattarai, Dongzhi Wang, Xiaochen He, Lihong Pan, John S. Clemmer, Lei Hou, Yingjie Chen

**Author notes:** **Correspondence:** Yingjie Chen, Ph.D.; or Lei Hou, Ph.D.

## Abstract

**BACKGROUND:** Interleukin-27 (IL-27) is a heterodimeric cytokine that serves as a bifunctional rheostat rather than an inherently pro- or anti-inflammatory signaling protein. However, the specific role of IL-27 in regulating systolic overload-induced cardiac inflammation and heart failure (HF) pathogenesis remains unknown.

**METHODS:** We investigated the effects of genetic IL-27 receptor deficiency (IL-27Rα knockout), pharmacological IL-27 blockade, and recombinant IL-27 administration on transverse aortic constriction (TAC)-induced HF in mice.

**RESULTS:** Cardiac IL-27 expression was significantly elevated in both murine and human HF tissues. The global genetic ablation of the IL-27 receptor (IL-27Rα) significantly suppressed TAC-induced cardiac inflammation, fibrosis, hypertrophy, HF progression, and mortality. Corroborating these protective effects, transcriptomic analysis (RNA-seq) revealed that IL-27Rα deficiency drastically suppressed pathways driving immune responses and antigen presentation, alongside the significant downregulation of networks governing systemic inflammation, pathogen infection, and extracellular matrix remodeling. Furthermore, pharmacological neutralization of IL-27 effectively attenuated TAC-induced left ventricular dysfunction, chamber dilation, myocardial hypertrophy, fibrosis, and leukocyte infiltration. Conversely, the administration of recombinant mouse IL-27 exacerbated the TAC-induced cardiac accumulation of multiple immune cell subsets, resulting in worsened cardiac fibrosis, cardiomyocyte hypertrophy, and overall HF progression.

**CONCLUSIONS:** Our findings demonstrate that IL-27 acts as a critical pathogenic driver of cardiac inflammation and HF development by modulating both cardiac immune cells (predominantly T cells) and non-immune cells, highlighting the IL-27 signaling axis as a promising therapeutic target.

## Introduction

Heart failure (HF) is a progressive, leading global cause of morbidity and mortality^1^. It is frequently driven by underlying conditions like coronary artery disease, hypertension, and autoimmunity, and is exacerbated by comorbidities such as diabetes and advanced age^2^. Chronic systolic overload—often secondary to hypertension or aortic stenosis—increases left ventricular (LV) wall stress and represents a primary risk factor for HF^3,4^. This sustained overload induces maladaptive remodeling, characterized by cardiomyocyte hypertrophy and fibrosis, ultimately culminating in HF^3,4^ alongside consequent pulmonary remodeling and right ventricular dysfunction^5,6^. Despite diverse etiologies, chronic low-grade inflammation remains a shared hallmark of HF^7–9^. Recent studies demonstrate that chronic inflammation actively promotes HF pathogenesis, and its targeted inhibition effectively attenuates disease progression in experimental models^7,8^. Clinically, IL-1β blockade with canakinumab significantly reduced major cardiovascular events in patients with prior myocardial infarction or active inflammation^10^, and dose-dependently decreased HF hospitalizations and related deaths^11^. However, the precise mechanisms by which inflammation drives HF remain incompletely understood.

Cardiac inflammation is driven by the dynamic interplay between innate and adaptive immunity^7,8^, a process orchestrated by cytokines, chemokines, and their receptors. Following initial cardiac injury, resident macrophages and recruited neutrophils initiate acute responses to clear cellular debris^8,12^. Subsequently, infiltrating pro-inflammatory monocytes and other innate immune cells produce inflammatory mediators such as TNF-α and IL-1β^13–15^. Persistent cardiac stress further drives maladaptive remodeling, triggering antigen-presenting cell (APC)-dependent T cell activation in lymphatic tissues and subsequent cardiac T cell infiltration^16–19^. Among these lymphocyte populations, CD8⁺ T cells contribute to HF progression in mice with preexisting LV dysfunction^20^.Studies by our group and others demonstrate that monocyte-derived macrophages^13^, CD8⁺ T cells^20^, dendritic cells^21^, NK1.1⁺ lymphocytes^22^, and other immune subsets actively drive the overload-induced inflammatory microenvironment and pathological remodeling. Correspondingly, pro-inflammatory cytokines (IL-1β TNF-α) exacerbate this overload-induced inflammation, fibrosis, and dysfunction, while their targeted inhibition or anti-inflammatory mediators (e.g., IL-10) offer cardioprotection^23^. In experimental models, inhibition of IL-1β signaling attenuated TAC-induced HF progression^15^, whereas IL-35 reduced pressure overload-induced cardiac inflammation and HF development^24^. Furthermore, we found that blocking TAC-induced T cell activation—whether via genetic deletion of CD28 or CD80/CD86, or by inhibiting APC-T cell interactions with CTLA4-Ig—significantly attenuates LV hypertrophy and HF^18,19^. Collectively, these findings highlight the essential pathogenic roles of macrophages, APCs, T cells, APC-T cell crosstalk, and pro-inflammatory cytokines in HF development.

The interleukin-27 (IL-27) signaling axis represents a critical immunoregulatory pathway that initiates potent pro-inflammatory responses during the early phases of infection^25–27^. Secreted primarily by activated antigen-presenting cells, the heterodimeric cytokine binds a receptor complex consisting of gp130 and the restricted IL-27Ra chain^28–30^, which is highly expressed on naive T cells, monocytes, macrophages, dendritic cells, and non-immune cells^27,29^. In immune cells, IL-27 receptor engagement activates JAK–STAT signaling, prominently involving STAT1 and STAT3, and can enhance IL-12 responsiveness in a cell context-dependent manner^27,31,32^. In naive CD4+ T cells, this axis directly upregulates the master transcription factor T-bet and the receptor subunit IL-12Rb2^31–33^, driving rapid differentiation and proliferation of Th1 effector cells^28,33^. Concurrently, IL-27 also enhances CD8⁺ T-cell function and promotes the generation of cytotoxic T lymphocytes^34^. Although macrophages, CD11c⁺ dendritic cells, and CD8⁺ T cells regulate pressure overload-induced cardiac inflammation and HF development^13,20,21^, the effect of IL-27 signaling axis on modulating systolic overload-induced cardiac inflammation and HF development is still unknown.

In the present study, we investigated the effects of IL-27 receptor knockout (IL-27Rα), IL-27 neutralizing antibodies, and the administration of recombinant IL-27 on transverse aortic constriction (TAC)-induced cardiac inflammation, cardiomyocyte hypertrophy, fibrosis, and left ventricular (LV) dysfunction in mice. We hypothesized that IL-27 signaling promotes pressure overload-induced HF by regulating myocardial inflammation and pathological remodeling. Using complementary genetic loss-of-function, pharmacological inhibition, and recombinant IL-27 gain-of-function approaches, we examined whether inhibition of IL-27 signaling attenuates, whereas exogenous IL-27 exacerbates, TAC-induced cardiac inflammation, remodeling, and dysfunction.

## Materials and Methods

### Experimental Mice and Protocols

Wild-type (WT) C57BL/6J mice (Strain #000664) and IL-27Rα KO (Strain #018078) mice were purchased from The Jackson Laboratory. Mice were housed in a temperature-controlled environment with a 12-hour light/dark cycle and provided ad libitum access to food and water. For all cohorts, samples were collected when apparent LV dysfunction was observed in at least one of the experimental groups following TAC. Samples were harvested for subsequent flow cytometric, histological, immunohistological, and biochemical analyses. All procedures were approved by the Institutional Animal Care and Use Committee at the University of Mississippi Medical Center.

### Pharmacological IL-27 Blockade and Recombinant IL-27 Administration

To evaluate the pharmacological inhibition of IL-27, WT female mice were subjected to TAC and randomly assigned to receive either an IL-27 neutralizing antibody (BioXCell, Catalog #BE0326) or an isotype control IgG. Treatments were administered via intraperitoneal injection at a dose of 250 µg/mouse every 3 days, starting 1-day post-TAC and continuing for a duration of 4 weeks. This dosage was adapted from a previous study^35^, as it was effectively suppressed the acute graft-versus-host disease in mice. Samples for histological analysis were collected 4 weeks post-TAC. For gain-of-function studies, WT mice received recombinant murine IL-27 (rmIL-27; BioLegend, #577406) via intraperitoneal injection at a dose of 2.5 µg/mouse/day, or a saline vehicle, beginning approximately 1 hour after TAC. The rmIL-27 dose was determined based on pilot dose-finding experiments with mild modification^36^.

### Flow Cytometry Analyses

The gating strategies of flow cytometry in different tissues are listed in the **Supplemental Figure 1 to Supplemental Figure 3**. *Detailed materials and methods are available in the online Data Supplement*.

## Results

### IL-27 expression is increased after HF

To investigate the inflammatory response associated with heart failure, the expression levels of specific cytokines were first evaluated in a wild type mice after TAC. TAC-induced pressure overload resulted in a significant elevation of cardiac IL-27, MCP1 (an adhesion molecule facilitating cardiac immune cell accumulation), and IL-1α protein levels compared to sham-operated controls (**Supplemental Fig 4**). Consistent with these quantitative findings, immunofluorescence staining of LV tissue demonstrated a substantial increase in the expression and structural distribution of the IL-27 subunit p28 (IL-27p28) in the failing hearts of TAC mice. To establish the clinical translational relevance of these observations, IL-27p28 expression was further examined in human heart samples. Mirroring the murine TAC model, LV tissue acquired from human patients with heart failure exhibited a highly significant upregulation of IL-27p28 compared to normal donor hearts, indicating that IL-27 signaling is a conserved feature of the cardiac remodeling process in both mice and humans.

### Genetic IL-27 inhibition by IL-27Ra KO significantly attenuated TAC-induced mortality and HF in mice of both sexes

To investigate the functional role of IL-27 signaling in the pathogenesis of pressure overload-induced heart failure, WT and IL-27Ra KO mice of both sexes were subjected to TAC. The experimental design incorporated baseline echocardiography followed by serial ultrasound monitoring at 2, 4, and 6 weeks post-surgery (**Figure 1A, left**). Strikingly, the genetic ablation of IL-27Ra conferred profound protection against TAC-induced mortality. As demonstrated by the Kaplan-Meier survival analysis (**Figure 1A, right**), WT controls exhibited a precipitous decline in survival following TAC, whereas IL-27Ra KO mice maintained a significantly higher survival rate (P=0.01).

**Fig 1:**
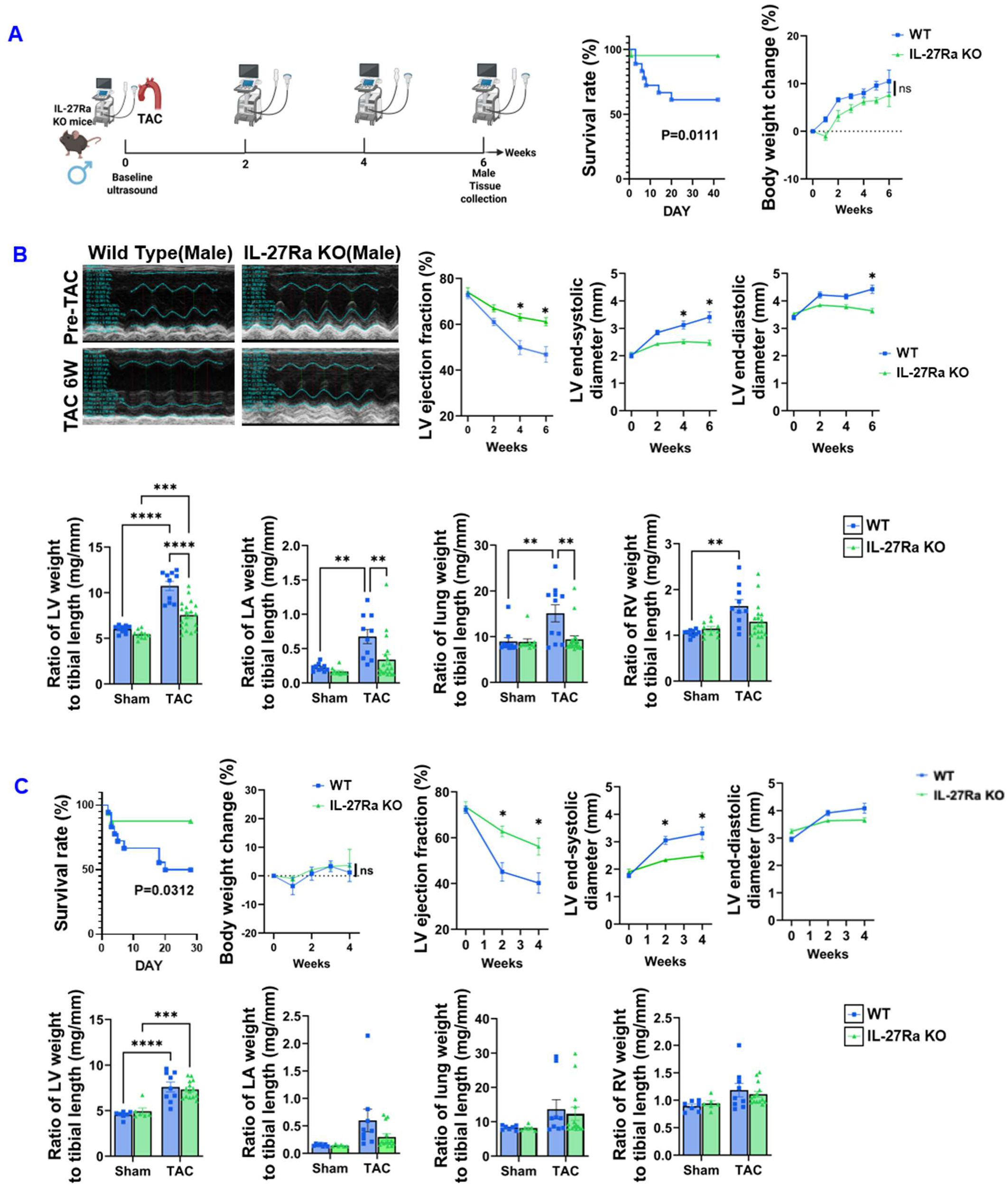
IL-27Ra KO protects against TAC-induced mortality and heart failure in male and female mice. **(A)** Schematic representation of the experimental timeline. WT and KO male mice underwent baseline echocardiography followed by TAC surgery, with subsequent ultrasound monitoring. Tissue collection occurred at 6 weeks for male cohorts. The accompanying Kaplan-Meier curve demonstrates a significantly higher survival rate in IL-27Ra KO mice (green) compared to WT mice (blue) following TAC (P=0.01). **(B)** Representative M-mode echocardiography images and quantitative time-course analysis of LVEF in male mice. KO mice maintained significantly higher LVEF post-TAC compared to the progressive decline observed in WT mice. Bar graphs quantify morphometric parameters, demonstrating reduced pathological remodeling in the KO group. **(C)** Corresponding survival, echocardiographic, and morphological assessments for the female cohort following TAC. Consistent with the initial cohort, IL-27Ra KO mice exhibited preserved cardiac function, improved survival, and reduced indices of cardiac hypertrophy and pulmonary congestion relative to WT controls. Data are visually represented comparing WT (blue) and IL-27Ra KO (green) groups.

Detailed echocardiographic assessments further elucidated the functional benefits of IL-27Ra deletion. Representative M-mode echocardiograms visually illustrated the preservation of left ventricular chamber dimensions and contractility in the KO cohort compared to the progressive dilation observed in WT mice (**Figure 1B, left**). Quantitative time-course analysis revealed that while WT mice developed severe heart failure characterized by a significant, progressive depression of left ventricular ejection fraction (LVEF), IL-27Ra KO mice exhibited remarkably preserved cardiac systolic function throughout the monitoring period (**Figure 1B, middle, Supplemental Table-2, Table-3**).

Post-mortem tissue morphometric analyses corroborated these functional findings; the bar graphs demonstrate that TAC-induced cardiac hypertrophy and pulmonary congestion—standard markers of pathological remodeling—were markedly blunted in the IL-27Ra KO cohorts (**Figure 1B, bottom, Supplemental Table-4, Supplemental Table-5**). Importantly, these robust protective effects were consistent across both sexes. Assessments in the complementary sex cohort mirrored these outcomes, revealing parallel improvements in survival, preserved LVEF, and reduced indices of pathological remodeling following TAC (**Figure 1C**). Together, these findings establish that intact IL-27 signaling is a critical mediator of adverse cardiac remodeling, and its genetic inhibition successfully attenuates heart failure progression irrespective of sex.

To evaluate the requirement of IL-27 signaling in pathological remodeling, we assessed cardiomyocyte hypertrophy in IL-27Ra KO mice following TAC. Male WT mice exhibited robust TAC-induced cardiomyocyte hypertrophy, which was significantly blunted in male IL-27Ra KO mice; conversely (**Supplemental Figure 5A, B**), females showed a milder overall hypertrophic response that remained unaltered by IL-27Ra deficiency (**Supplemental Figure 5C**).

### Transcriptomic profiling reveals attenuation of profibrotic gene networks and cardiac fibrosis in IL-27Ra KO mice

To delineate the downstream molecular mechanisms by which IL-27Ra deletion protects against pressure overload, bulk RNA-sequencing was performed on LV tissues from female WT and IL-27Ra KO mice. Principal component analysis (PCA) and hierarchical clustering of the transcriptomic data demonstrated highly distinct gene expression profiles among the groups. Notably, the WT TAC group clustered entirely separately from the sham controls, whereas the KO TAC group exhibited an intermediate transcriptional signature that remained much closer to the baseline control state (**Figure 2A**). Differential gene expression analysis comparing KO TAC to WT TAC tissues, visualized via volcano plot, revealed a widespread downregulation of numerous pathological and profibrotic transcripts in the KO group, prominently including genes such as *Postn*, *Cilp*, *Col12a1*, *Col8a2*, and *Myh7* (**Figure 2B**). Gene Ontology (GO) analyses of these differentially expressed genes highlighted a dominant representation of terms associated with the extracellular matrix (ECM), including “collagen-containing extracellular matrix,” “basement membrane,” and “collagen trimer” organization (**Figure 2C**).

**Figure 2:**
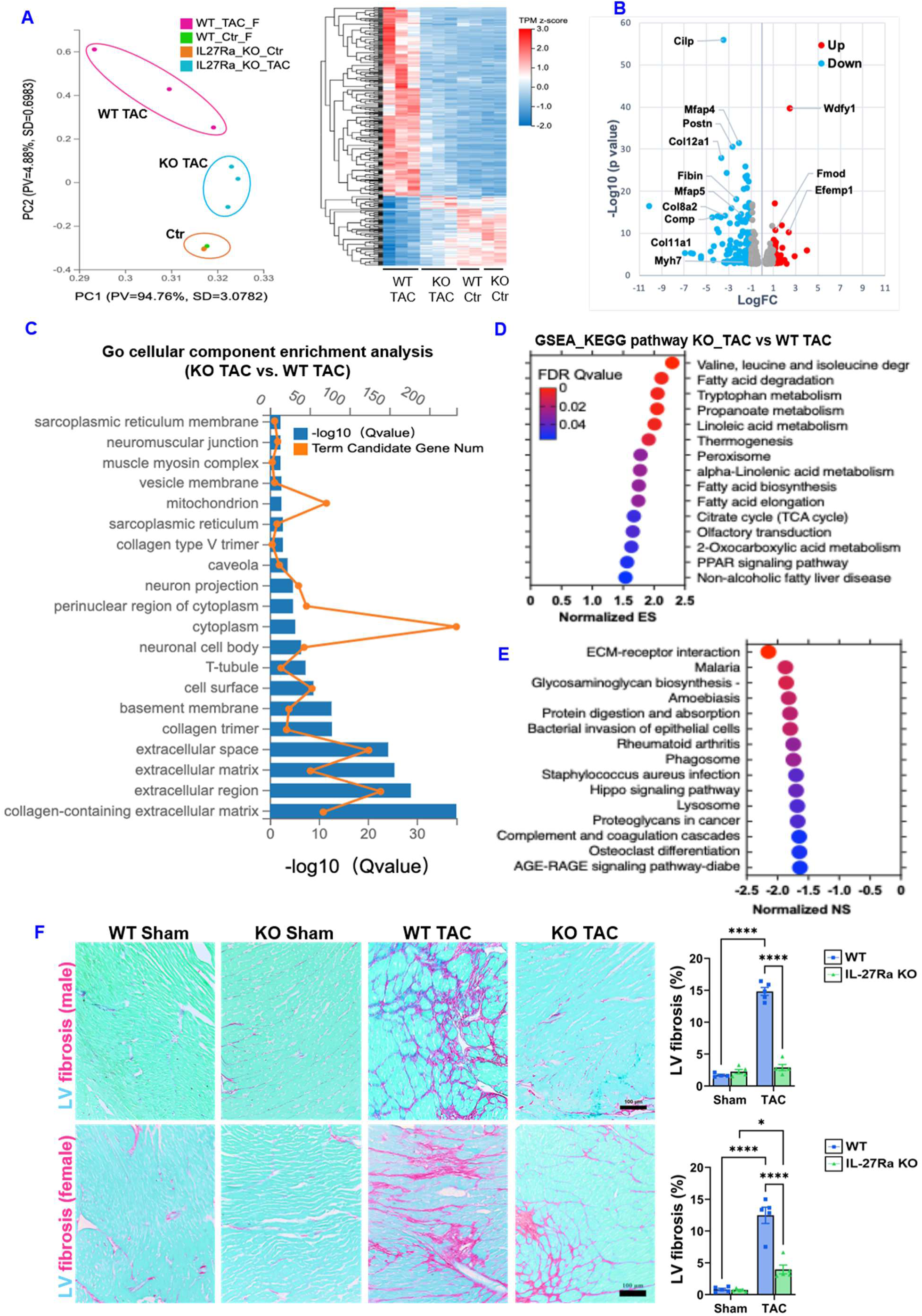
IL-27Ra deletion blunts fibrotic gene transcription and mitigates TAC-induced cardiac fibrosis. **(A)** Principal component analysis (PCA) plot and hierarchical clustering heatmap of bulk RNA-seq data from female WT and IL-27Ra KO LV tissues (Sham vs. TAC), demonstrating the global transcriptomic shift and intermediate phenotype of the KO TAC group. **(B)** Volcano plot illustrating differentially expressed genes between KO TAC and WT TAC tissues. Blue dots denote significantly downregulated genes in the KO group (e.g., *Postn*, *Cilp*, *Col12a1*, *Myh7*), while red dots indicate upregulated genes. **(C)** GO enrichment analysis displaying the top cellular component terms downregulated in the KO TAC vs. WT TAC comparison, highlighting a strong association with the extracellular matrix and collagen structures. **(D, E)** Pathway enrichment dot plots further supporting the suppression of structural and fibrotic remodeling networks in the KO cohort. **(F)** Representative histological images (left) of LV tissue from male and female mice stained for fibrosis (collagen deposition in pink/red) following TAC. Accompanying bar graphs (right) quantify the percentage of fibrotic area, showing a significant reduction in fibrosis in both male and female IL-27Ra KO mice compared to WT controls.

To identify the core biological pathways altered in this model, we performed pathway enrichment analysis (**Figure 2D**). The analysis revealed a distinct dichotomy between upregulated and downregulated gene sets. Pathways exhibiting positive enrichment (Normalized ES > 0) were overwhelmingly associated with mitochondrial and cellular metabolism, with the most significant upregulation seen in amino acid degradation (valine, leucine, and isoleucine), fatty acid degradation, and the citrate (TCA) cycle (**Figure 2D, Supplemental Figure 6A,B**). Conversely, pathways exhibiting negative enrichment scores (Normalized NS < 0) were heavily dominated by ECM remodeling, immune and inflammatory responses, and infectious disease pathways (**Figure 2E, Supplemental Figure 6C**). Notably, ECM-receptor interaction and various pathogen-response pathways showed the most significant downregulation, highlighting a broad suppression of specific structural and inflammatory signaling cascades alongside the enhancement of metabolic function.

Consistent with this transcriptomic suppression of ECM and collagen-related pathways, histological evaluation of LV tissue confirmed a striking reduction in reactive cardiac fibrosis in the IL-27Ra KO mice. Morphological staining of heart sections revealed that while WT mice developed severe, widespread interstitial fibrosis following TAC—evidenced by extensive collagen deposition (stained in pink/red)—this structural remodeling was robustly attenuated in the IL-27Ra KO cohorts (**Figure 2F, left**). Quantitative analysis of the fibrotic area corroborated these visual findings, demonstrating a highly significant reduction in LV fibrosis in both male and female IL-27Ra KO mice compared to their WT counterparts subjected to TAC (**Figure 2F, right**). Together, these molecular and histological data indicate that IL-27Ra ablation preserves cardiac function by halting the severe fibrotic remodeling cascade induced by chronic TAC.

### Identification of significantly changed genes associated with immunity and inflammation in IL-27Ra KO mice

Functional classification of differentially expressed genes revealed a robust enrichment in pathways related to the “Immune system” and infectious diseases (viral, bacterial, and parasitic) (**Supplemental Figure 7A**). Consistent with this functional classification, hierarchical clustering and expression heatmap analysis identified a distinct module of highly upregulated genes intricately linked to immune activation, inflammation, and pathogen response (**Supplemental Figure 7B**). Among the top upregulated transcripts in this dominant cluster were several critical components of the complement system (***C4b***, ***C7***, ***Cfh***, and ***Serping1***), which play a vital role in innate immunity and inflammation. Furthermore, the analysis revealed prominent upregulation of ***Lbp*** (Lipopolysaccharide binding protein), a classic acute-phase protein involved in the immune response to bacterial infection; ***Spp1*** (Osteopontin), a key pro-inflammatory and pro-fibrotic cytokine; the chemokine ***Ccl21a***, which directs immune cell migration; and ***Cd44***, a critical surface receptor mediating leukocyte activation and adhesion. Together, these transcriptomic signatures confirm a widespread and potent upregulation of networks governing innate immunity and inflammatory signaling.

### IL-27Ra Deletion Attenuates Global Leukocyte Infiltration and Modulates Myocardial Immune Cell Composition Following TAC

To characterize the inflammatory cellular response during cardiac remodeling, general leukocyte infiltration in the LV was evaluated using CD45 immunofluorescence staining. As expected, pressure overload via TAC provoked a robust inflammatory response in WT mice, characterized by a massive accumulation of CD45-positive immune cells within the myocardium. In stark contrast, genetic ablation of IL-27Ra significantly blunted this TAC-induced leukocyte infiltration. Importantly, quantitative analysis confirmed that this marked reduction in global myocardial inflammation was consistent across both male and female cohorts (**Figure 3A**), indicating a sex-independent mechanism by which IL-27 signaling promotes widespread immune cell recruitment to the stressed heart.

**Figure 3:**
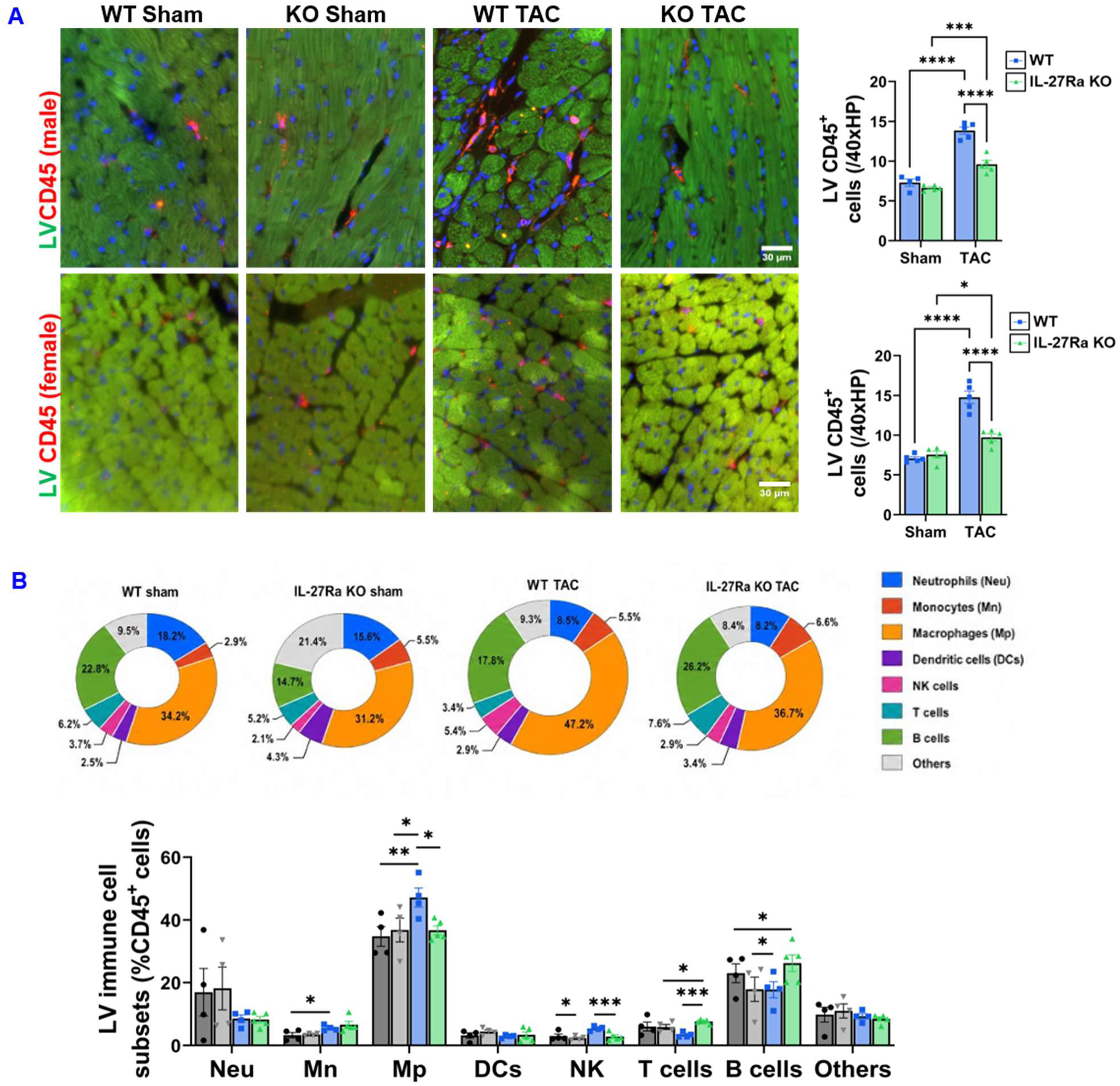
IL-27Ra deficiency blunts TAC-induced myocardial leukocyte infiltration and alters immune cell subset composition. **(A)** Representative immunofluorescence images and corresponding quantification of CD45+ leukocytes (red) in the left ventricle (LV) of male (top panel) and female (bottom panel) mice. Genetic deletion of IL-27Ra significantly reduced the total accumulation of CD45+ immune cells following TAC in both sexes. **(B)** Flow cytometric profiling of LV immune cell subsets in male mice. The donut charts visually map the shifting proportions of specific leukocyte lineages within the total CD45+ population across the WT and IL-27Ra KO sham and TAC groups. The accompanying grouped bar graph quantifies the relative percentages of neutrophils (Neu), monocytes (Mn), macrophages (Mp), dendritic cells (DCs), natural killer (NK) cells, T cells, and B cells, highlighting a significant attenuation of macrophage expansion in the IL-27Ra KO TAC cohort. Statistical significance between relevant groups is denoted by asterisks.

To further dissect the specific leukocyte subpopulations driving this inflammatory response, flow cytometric profiling of the LV was performed on male mice. Comprehensive analysis of the total CD45^+^ leukocyte compartment revealed dynamic, TAC-induced shifts in the immune cell landscape. Macrophages constituted the most dominant immune cell population in the remodeling heart and underwent a highly significant expansion in WT mice following TAC. However, consistent with the overall reduction in inflammation, this pressure-driven macrophage expansion was significantly attenuated in the IL-27Ra KO mice (**Figure 3B**). Furthermore, IL-27Ra deletion altered the composition of other immune subsets; for example, the relative proportions of T cells, B cells, and NK cells within the CD45^+^ pool were significantly higher or preserved in the KO TAC cohort compared to the WT TAC controls. Collectively, these data suggest that intact IL-27 signaling is critical not only for driving total leukocyte recruitment but also for orchestrating the specific, macrophage-heavy inflammatory microenvironment typical of heart failure.

### Pharmacological inhibition of IL-27 signaling significantly attenuated TAC-induced cardiac hypertrophy, dysfunction, and immune cell accumulation in wild type mice

To investigate the therapeutic potential of IL-27 blockade in pressure overload-induced heart failure, mice subjected to TAC were treated with an IL-27 neutralizing monoclonal antibody (IL-27 mAb; 250 µg/mouse, every 3 days i.p.) or an IgG isotype control over a 4-week period. Echocardiographic analysis over the disease course revealed that systemic IL-27 blockade significantly preserved left ventricular systolic function and attenuated progressive chamber dilation compared to the IgG-treated TAC cohort (**Figure 4A, Supplemental Table-6**). Consistent with these functional improvements, post-mortem gravimetric analysis at 4 weeks post-TAC demonstrated that IL-27 mAb administration significantly blunted pathological cardiac hypertrophy and indices of pulmonary congestion, as evidenced by broadly reduced normalized organ weights (**Figure 4B, Supplemental Table-7**). Furthermore, histological evaluation utilizing Sirius Red staining and immunofluorescence targeting CD45 showed that neutralizing IL-27 markedly suppressed TAC-induced adverse myocardial remodeling, significantly reducing both interstitial fibrosis and total leukocyte infiltration within the ventricular myocardium relative to IgG controls (**Figure 4C**).

**Figure 4.**
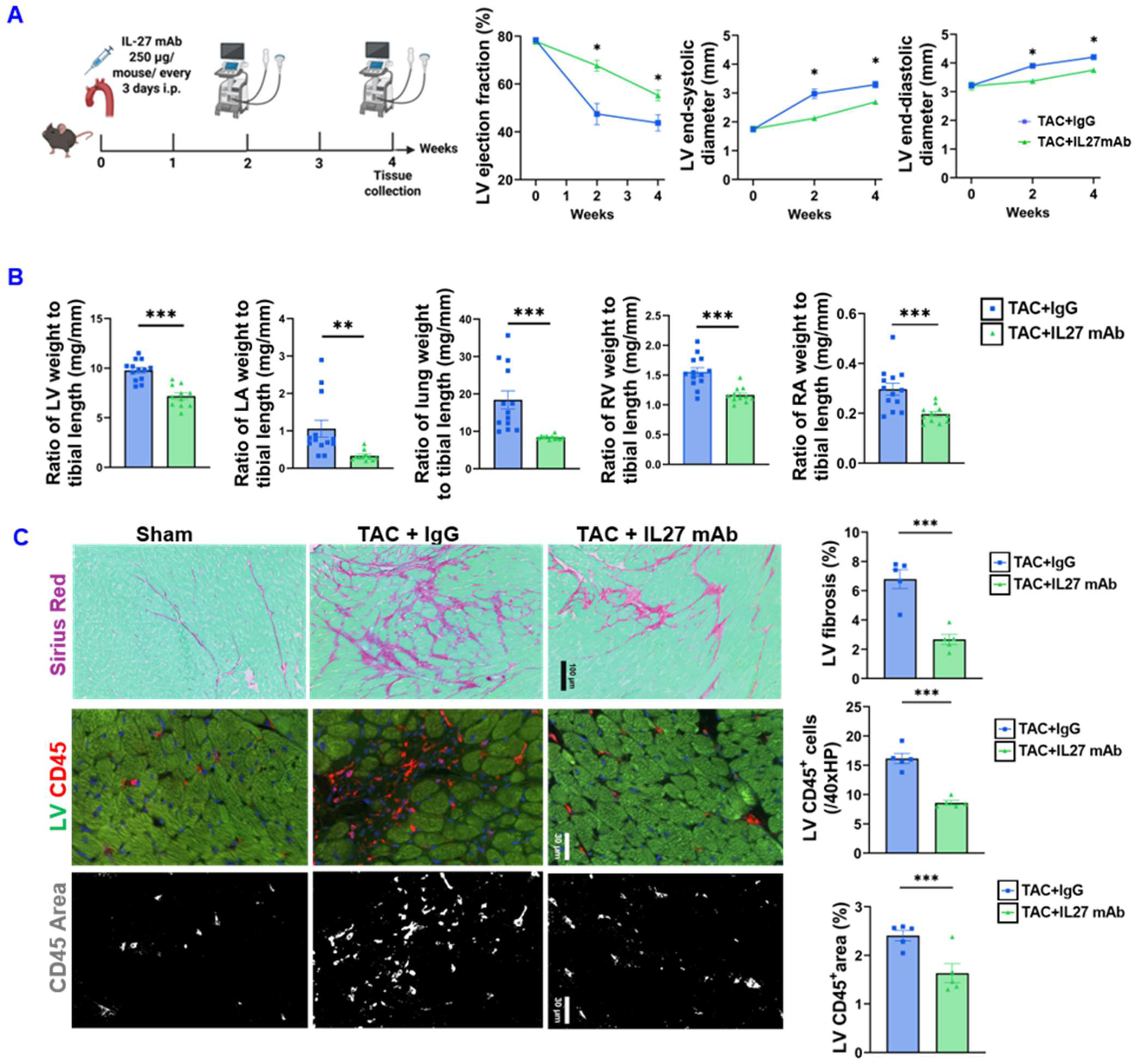
IL-27 neutralizing antibody attenuates TAC-induced cardiac dysfunction, hypertrophy, and myocardial inflammation. **(A)** Experimental schematic detailing the transverse aortic constriction (TAC) model, followed by the intraperitoneal (i.p.) administration of either an IL-27 monoclonal antibody (IL-27 mAb; 250 µg/mouse every 3 days) or an IgG vehicle control for 4 weeks. Accompanying line graphs illustrate longitudinal echocardiographic assessments of left ventricular function and chamber dimensions at 2 and 4 weeks post-surgery. **(B)** Post-mortem gravimetric analysis at 4 weeks post-TAC. Quantitative bar graphs represent normalized organ weights, comparing the extent of cardiac hypertrophy and congestion between the TAC+IgG and TAC+IL27 mAb cohorts. **(C)** Representative histological and immunofluorescence images evaluating myocardial tissue remodeling. Sirius Red staining (top row) assesses interstitial fibrosis, while LV CD45 staining (middle row) and corresponding isolated mask areas (bottom row) evaluate leukocyte infiltration. Accompanying bar graphs quantify the reduction in fibrotic area and CD45+ cellular accumulation in the IL-27 mAb treated group compared to IgG controls.

### IL-27 administration significantly exacerbated TAC-induced cardiac hypertrophy, dysfunction, fibrosis, and immune cell accumulation in wild type mice

To investigate the in vivo effects of Interleukin-27 on pathological cardiac remodeling, wild type mice subjected to TAC were administered daily evening intraperitoneal injections of either recombinant mouse IL-27 (rmIL-27; 2.5 µg/mouse) or a saline vehicle control over a two-week period. Echocardiographic analysis revealed that rmIL-27 administration significantly accelerated the deterioration of cardiac function, evidenced by a marked reduction in LV ejection fraction and a concomitant increase in LV end-systolic diameter at both one and two weeks post-TAC compared to saline-treated controls, while LV end-diastolic diameter remained unaltered (**Figure 5A, Supplemental Table-8**). Consistent with the functional decline observed via echocardiography, post-mortem gravimetric analysis at two weeks demonstrated that rmIL-27 treatment significantly exacerbated TAC-induced cardiac hypertrophy; specifically, the ratios of whole heart, left ventricle, left atrium, and right ventricle weights normalized to tibial length were all significantly elevated in the TAC+rmIL27 cohort (**Figure 5B, Supplemental Table-9**). Furthermore, the rmIL-27 treated mice exhibited a significantly increased ratio of lung weight to body weight, indicative of worsened pulmonary congestion secondary to progressing heart failure, whereas right atrial weight parameters showed no significant difference between the experimental groups. Histological evaluation and immunofluorescence staining revealed that while TAC induced significant interstitial fibrosis and CD45-positive leukocyte infiltration compared to sham-operated controls, administration of rmIL-27 profoundly exacerbated both fibrotic remodeling and total CD45+ cell accumulation (**Figure 5C**).

**Figure 5.**
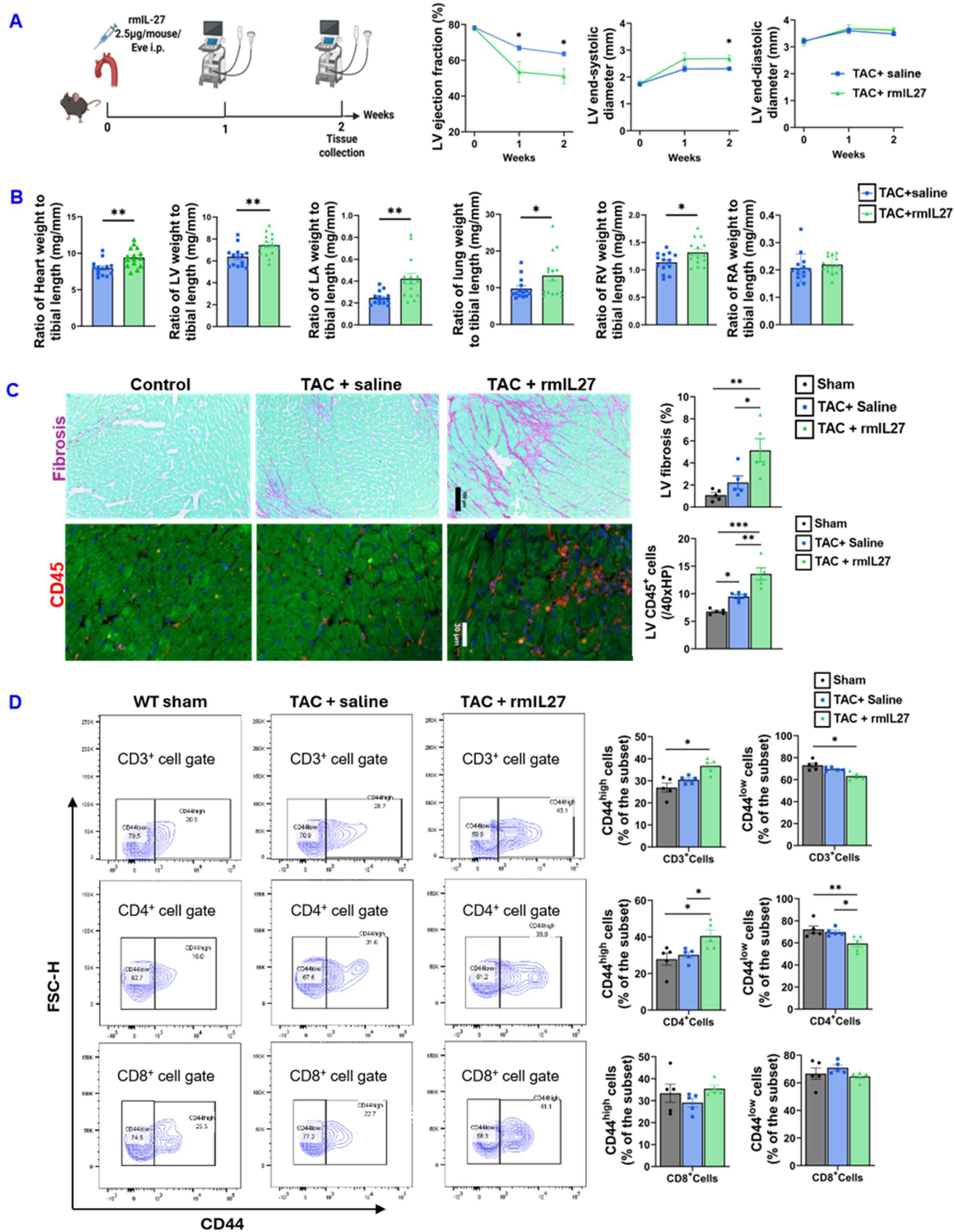
IL-27 administration exacerbates TAC-induced cardiac hypertrophy, dysfunction, and myocardial inflammation. **(A)** Experimental schematic and echocardiographic quantification of left ventricular (LV) ejection fraction, end-systolic diameter, and end-diastolic diameter in mice treated with daily rmIL-27 (2.5 µg/mouse) or saline for 2 weeks post-TAC. **(B)** Post-mortem gravimetric analysis of cardiac remodeling and pulmonary congestion, showing normalized weights for the heart, LV, left atrium, right ventricle, right atrium, and lungs. **(C)** Representative histological and immunofluorescence images with corresponding quantification assessing myocardial fibrosis and total CD45+ leukocyte infiltration. **(D)** Flow cytometric analysis of T cell activation, demonstrating the selective upregulation of the activation marker CD44 within the CD4+ helper T cell compartment following rmIL-27 administration. Data are represented as mean +/-SEM. Statistical significance between the TAC+saline and TAC+rmIL27 groups is denoted by * p < 0.05 and **p < 0.01.

### IL-27 administration significantly exacerbated TAC-induced LV multiple immune cell accumulation, and T cell activation in wild type mice

To further characterize the impact of IL-27 on pathological cardiac remodeling, we assessed myocardial immune cell subsets within the ventricular tissue (**Figure 5D–E, Supplemental Figure 8**). Detailed profiling of these infiltrating immune cell populations demonstrated a predominantly macrophage-driven inflammatory response—accounting for over 50% of the detected immune cells across all TAC groups—alongside notable populations of B cells, T cells, and neutrophils; however, the relative proportions of these specific immune subsets remained largely comparable between the saline- and rmIL-27-treated TAC cohorts (**Supplemental Figure 8**). Additional quantitative assessments confirmed that while TAC surgery significantly elevated specific pathological cell counts or markers relative to sham controls, these secondary parameters reached similar plateau levels in both TAC groups regardless of rmIL-27 treatment (**Supplemental Figure 8**), suggesting that IL-27 drives specific fibrotic and total leukocyte exacerbations rather than shifting the proportional composition of the immune infiltrate.

Our previous studies showed that cardiac T cell activation contribute to TAC-induced cardiac inflammation and HF development. To determine the specific T cell populations activated by IL-27 during pathological cardiac remodeling, we evaluated the expression of the activation marker CD44 across distinct T cell subsets using flow cytometry (**Figure 5D**). Analysis of the total T cell pool (CD3^+^ gate) revealed that rmIL-27 administration following TAC significantly increased the proportion of activated CD44^high^ cells and reciprocally decreased the naive CD44^low^ population compared to sham controls. Further subtyping demonstrated that this activation shift was predominantly driven by the helper T cell compartment; specifically, the TAC + rmIL27 cohort exhibited a statistically significant elevation in CD44^high^ CD4^+^ T cells, and a corresponding reduction in CD44^low^ CD4^+^ T cells, when compared to both the WT sham and TAC + saline vehicle groups. In contrast, the activation status of cytotoxic T cells (CD8^+^ gate) remained largely unaltered, with no significant differences observed in CD44 expression across any of the experimental conditions, indicating that IL-27 administration selectively enhances CD4^+^ T cell activation in the context of pressure overload.

### Impact of TAC and IL-27 administration on immune cell composition in the draining lymph nodes in wild type mice

To further characterize the regional inflammatory response to pressure overload and exogenous IL-27, we evaluated the draining lymph nodes associated with the pulmonary and cardiac tissues (**Supplemental Figure 9**). Macroscopic examination revealed marked lymph node enlargement (lymphadenopathy) in mice subjected to TAC compared to sham-operated controls (**Supplemental Figure 9A**), indicative of a robust regional immune activation. Despite this overall expansion in lymph node size, flow cytometric profiling demonstrated that the relative compositional breakdown of the resident immune cells remained remarkably conserved across all experimental groups. Specifically, the lymph nodes were consistently dominated by B cells (∼40–42%), CD4^+^ T cells (∼26–30%), and CD8^+^ T cells (∼24–26%), with only minor fluctuations in γδT cells, dendritic cells, and NK cells **Supplemental Figure 9B,C**. Subsequent quantitative analysis of these individual leukocyte subsets confirmed that while TAC drives overall regional lymphatic expansion, systemic administration of rmIL-27 does not significantly skew or alter the proportional distribution of the primary immune cell populations within the draining lymph nodes when compared to saline-treated TAC controls.

Having observed an IL-27-mediated activation of T cells within the myocardium, we next sought to determine if this inflammatory signaling influenced T cell activation within the regional lymphoid tissues (**Supplemental Figure 10**). Flow cytometric analysis of the draining lymph nodes was performed to evaluate the expression of the activation marker CD44 across distinct T cell subsets. Within the total T cell pool (CD3^+^ gate), mice subjected to transverse aortic constriction (TAC) and treated with rmIL-27 exhibited a significant increase in the proportion of activated CD44^high^ cells, alongside a reciprocal decrease in naive CD44^low^ cells, compared to WT sham controls. Consistent with the shifts observed in the cardiac infiltrate, this robust transition towards an activated phenotype was predominantly localized to the helper T cell compartment. Specifically, the TAC + rmIL27 cohort demonstrated a statistically significant elevation in the percentage of CD44^high^ CD4^+^ T cells, and a corresponding reduction in CD44^low^ cells, relative to the sham group. In contrast, the activation status of cytotoxic T cells (CD8^+^ gate) remained largely stable, with no statistically significant differences in CD44 expression observed across the experimental groups. These findings indicate that systemic IL-27 administration actively augments regional CD4^+^ T cell activation within the draining lymph nodes during pressure overload.

### Genetic IL-27 inhibition by IL-27Ra KO significantly attenuated TAC-induced pulmonary inflammation and fibrosis in mice of both sexes

Because left-sided heart failure following TAC frequently leads to secondary pulmonary congestion and pathological remodeling, we next evaluated lung tissue for inflammation and fibrosis (**Supplemental Figure 11**). Immunofluorescence staining for the pan-leukocyte marker CD45 revealed that TAC surgery induced a marked accumulation of immune cells within the lungs of both male and female WT mice compared to sham-operated controls. However, this secondary pulmonary inflammatory response was significantly attenuated in both male and female IL-27Ra KO mice (**Supplemental Figure 11A**). Furthermore, histological assessment demonstrated a significant increase in interstitial lung fibrosis in WT mice subjected to TAC. Consistent with the blunted immune infiltration, IL-27Ra deficiency profoundly protected against TAC-induced pulmonary fibrotic remodeling in both sexes, maintaining fibrotic areas at or near baseline sham levels (**Supplemental Figure 11B**). Together, these findings indicate that intact IL-27 receptor signaling contributes heavily to the secondary pulmonary inflammation and fibrosis associated with pressure overload-induced heart failure.

To evaluate secondary pulmonary vascular remodeling, we further assessed lung smooth muscle alpha-actin (α-SMA) expression in male mice (**Supplemental Figure 12**). Immunofluorescence revealed that TAC induced significant pulmonary vascular muscularization—characterized by increased perivascular α-SMA deposition—in WT mice compared to sham controls. Importantly, this pathological remodeling was profoundly blunted in IL-27Ra KO mice. These data indicate that IL-27 receptor signaling drives secondary pulmonary vascular remodeling during pressure overload.

### Pharmacological inhibition of IL-27 signaling significantly attenuated TAC-induced secondary pulmonary inflammation, and macrophage accumulation

To investigate the therapeutic efficacy of IL-27 neutralization on secondary pulmonary inflammation, we comprehensively profiled the lung immune cell infiltrate using flow cytometry. Following TAC, IgG-treated mice exhibited a dramatic shift in the pulmonary immune landscape, driven predominantly by a massive accumulation of macrophages, which expanded to represent 38.1% of the immune compartment compared to just 21.8% in sham controls (**Supplemental Figure 13A, B**). Conversely, systemic administration of an IL27 mAb significantly blunted this pathological macrophage expansion, reducing their proportion to 24.5%. This reduction in macrophage burden coincided with the relative preservation of other resident lymphocytes, including T cells, dendritic cells (DCs), and B cells within the pulmonary tissue. Further quantitative analysis of specific immune cell subsets confirmed that IL-27 blockade broadly suppressed TAC-induced inflammatory shifts, normalizing absolute cell counts and specific activation parameters compared to the IgG vehicle cohort (**Supplemental Figure 13C-E**). These findings demonstrate that pharmacological inhibition of IL-27 effectively protects against secondary pulmonary inflammation and pathological macrophage infiltration during pressure overload-induced heart failure.

### Pharmacological inhibition of IL-27 signaling significantly attenuated TAC-induced secondary pulmonary activation of macrophages, dendritic cells, and T cells

Because pulmonary macrophages and dendritic cells are recognized as professional APCs that play a crucial role in shaping local inflammatory responses, we next investigated their activation status by evaluating Major Histocompatibility Complex class II (MHC-II) expression (**Supplemental Figure 14**). Following TAC, IgG-treated mice exhibited a striking upregulation of MHC-II across multiple pulmonary myeloid populations. This robust pro-inflammatory activation was evident in alveolar macrophages (Alvo-Mp), monocyte-derived macrophages (Mo-Mp), interstitial macrophages (Int-Mp), and CD11c^+^ dendritic cells), all of which showed a substantial increase in the proportion of MHC-II ^high^ cells compared to sham controls. Strikingly, systemic administration of the IL-27 neutralizing antibody completely abrogated this TAC-induced upregulation (**Supplemental Figure 14**). IL-27 blockade profoundly suppressed MHC-II expression across all assessed macrophage subpopulations and dendritic cells, effectively maintaining their activation status at or near baseline sham levels (**Supplemental Figure 14**). These data demonstrate that pharmacological inhibition of IL-27 broadly blunts the pro-inflammatory activation and antigen-presenting capacity of the pulmonary myeloid compartment during pressure overload.

To determine whether IL-27 drives pulmonary T cell activation during secondary lung inflammation, we analyzed CD44 expression on T cells isolated from the lung tissue. Following TAC, IgG-treated mice exhibited a massive increase in the proportion of activated (CD44^high^) T cells, with a reciprocal decrease in naive/resting (CD44^low^) T cells. This robust activation was observed across the total (CD3^+^), helper (CD4^+^), and cytotoxic (CD8^+^) T cell compartments (**Supplemental Figure 15**). Importantly, systemic treatment with an IL-27 neutralizing antibody significantly blunted this TAC-induced activation, profoundly suppressing CD44 expression and restoring the naive T cell profile across all assessed subsets (**Supplemental Figure 15**). These data demonstrate that pharmacological blockade of IL-27 effectively prevents secondary pulmonary T cell activation following pressure overload.

## Discussion

While IL-27 is generally recognized as a heterodimeric cytokine that serves as a bifunctional rheostat rather than an inherently pro- or anti-inflammatory signaling protein, in the present study, we provide comprehensive evidence establishing the cytokine IL-27 as a critical pathogenic driver of pressure overload–induced HF. This investigation was prompted by our observation that IL-27p28 expression was increased in myocardial samples from patients with end-stage ischemic cardiomyopathy, underscoring its immediate translational relevance. Utilizing a rigorous, multipronged approach—incorporating genetic ablation (IL-27Ra KO), pharmacological inhibition (neutralizing antibody), and gain-of-function (recombinant IL-27) models—we definitively demonstrate that IL-27 signaling is requisite for maladaptive cardiac remodeling and dysfunction. Mechanistically, our major new findings reveal that IL-27 actively promotes the cardiac infiltration of inflammatory myeloid and lymphoid cells, orchestrates pathological fibrosis via extracellular matrix reorganization, and dramatically exacerbates pulmonary vascular remodeling secondary to LV failure. Together, these data fundamentally challenge the prevailing paradigm of IL-27 strictly as an immunoregulatory molecule, redefining it instead as a potent therapeutic target in sterile cardiac inflammation.

Our findings position IL-27 as a convergent effector within the broader IL-12 cytokine family, extending our previous observations regarding the IL-12/IL-23 axis^37^. We previously demonstrated that the shared p40 subunit (IL-12β) of IL-12 and IL-23 is essential for HF progression^38,39^. IL-27, a heterodimer of p28 and EBI3 (a p40-related protein), shares distinct structural homology and downstream signaling pathways (e.g., JAK-STAT) with IL-12 and IL-23^28,29,40^. While some studies suggest compensatory relationships among family members, the data presented here indicate that in the context of pressure overload, these cytokines do not counterbalance each other but rather act synergistically to sustain a deleterious Type 1–biased inflammatory microenvironment. This distinction is crucial; because clinical strategies targeting the p40 subunit (e.g., ustekinumab) are already in use, our data suggests that isolated blockade of IL-27 or its receptor may offer complementary or even superior efficacy in cardiovascular disease.

A striking aspect of our study is the unambiguous pro-inflammatory phenotype of IL-27 in the failing heart, which sharply contrasts with its well-characterized immunoregulatory functions in autoimmunity and infection—such as suppressing Th17 responses or inducing IL-10^27,29^. This discrepancy emphasizes the highly context-dependent nature of cytokine signaling^27,29^. Consistent with this context dependence, IL-27R signaling has been reported to limit atherosclerosis in Ldlr^−/−^ mice^41^, whereas our findings indicate a predominantly pathogenic role during pressure overload-induced HF. In the setting of sterile cardiac injury, unlike in pathogen-driven responses, the “regulatory” brake of IL-27 appears completely absent or overwhelmed. Instead, our gain-of-function data show that IL-27 actively exacerbates T cell activation (CD44+) and pathological macrophage recruitment. This supports emerging evidence in other sterile inflammatory conditions where IL-27 acts as a pro-inflammatory amplifier, suggesting that the cytokine’s ultimate function is dictated by the specific inflammatory cues of the local tissue microenvironment.

Beyond immune modulation, our transcriptomic analysis reveals a profound role for IL-27 in regulating cardiac fibrosis. IL-27Ra deficiency resulted in a global normalization of gene programs governing ECM organization and collagen fibril assembly (e.g., *Col1a1*, *Postn*), which correlated strongly with reduced interstitial fibrosis. Current antifibrotic strategies often focus on the classical TGF-β signaling pathway^42,43^; however, clinical outcomes targeting this axis have been highly variable. Our data identifies IL-27 as a potent, inflammation-driven pro-fibrotic signal that may function distinctly from, or synergistically with, these classical pathways. Although IL-27 is known to signal via STAT1 and STAT3 in immune cells^27,29^, its robust regulation of the cardiac ECM implies a critical immune-fibrotic crosstalk. Our results suggest that this pro-fibrotic effect is likely dual-pronged: operating indirectly via the recruitment of pro-fibrotic macrophages and T cells^13,44^, and potentially through direct modulation of the resident fibroblast phenotype—a hypothesis that warrants future cell-specific mechanistic investigation^45,46^.

Importantly, our study highlights the efficacy of IL-27 blockade in mitigating the transition from LV failure to pulmonary hypertension and right ventricular hypertrophy. Pulmonary remodeling is a hallmark of advanced HF and a primary determinant of mortality^5,6,47^. We observed that IL-27 inhibition significantly reduced pulmonary vascular muscularization and local inflammation. This protection is likely twofold: providing a hemodynamic benefit from preserved LV function (thereby reducing backward pressure) alongside a direct anti-inflammatory effect in the pulmonary bed, as evidenced by the notably reduced antigen-presenting capacity (MHC-II expression) of pulmonary macrophages. This positions IL-27 inhibition as a comprehensive strategy that not only protects the myocardium but also preserves the pulmonary vascular bed, potentially delaying the onset of Group 2 Pulmonary Hypertension.

From a translational perspective, the profound efficacy of the anti-IL-27 neutralizing antibody in our model provides a compelling proof-of-concept for therapeutic intervention. Compared with broad immunosuppression, targeting a defined cytokine pathway may provide a more selective approach to modulating chronic inflammation in HF^10,11^. In conclusion, we have identified IL-27 as a critical, maladaptive checkpoint in the progression of heart failure. Targeting this specific pathway offers a novel and viable clinical avenue to suppress pathological inflammation and fibrosis, preserving both cardiac and pulmonary function in the setting of chronic pressure overload.

## Notes

Conflict of Interest: None.

### Competing Interest Statement

The authors have declared no competing interest.

